# Learning Discrete Cell and Niche Codes from Spatial Transcriptomics Using Dual Residual Vector Quantization

**DOI:** 10.64898/2026.08.07.743490

**Authors:** Sebastian Birk, Arpit Merchant, Amirhossein Vahidi, Fabian J. Theis, Mo Lotfollahi

**Author notes:** Joint last authors (Fabian J. Theis, Mo Lotfollahi).

## Abstract

Spatially-resolved transcriptomics (SRT) measures gene expression at single-cell resolution while preserving each cell’s spatial location, enabling the joint study of *cell identity* and cellular *niche*, the recurring microenvironment that organises tissue function. Existing representation-learning methods typically capture only one of these axes at a time. We present SQUINT, a graph vector-quantized variational autoencoder (VQ-VAE) that learns *two* disjoint codebooks per cell from a shared architecture: a *cell codebook* quantising the per-cell embedding before neighbourhood aggregation, biased toward cell-intrinsic identity, and a *niche codebook* quantising the embedding after graph neural network (GNN) aggregation, biased toward spatial context. Both use residual vector quantization, giving a coarse-to-fine discrete-token hierarchy. SQUINT is trained with per-branch negative-binomial reconstruction objectives and three domain-motivated components that we show are crucial: a within-section cosine adjacency loss that anchors the niche codes in the spatial graph, a cross-section contrastive loss on the cell latents that aligns transcriptomically matched cells, and a decoder section covariate that absorbs batch effects. Across three datasets spanning four spatial assays (STARmap, MERFISH, CosMx, Xenium) and four tasks – niche identification, cell-type identification, cross-section integration, and spatial gene-expression imputation in held-out regions – SQUINT outperforms or is competitive with strong baselines on identification and achieves the most faithful cross-section integration. The resulting discrete vocabulary makes tissues directly consumable by transformer-style foundation models and enables one-step query-to-reference atlas mapping via code-distribution similarity, which we demonstrate on a CosMx human non-small-cell lung cancer cohort.

## 1. Introduction

Spatially-resolved transcriptomics (SRT) platforms now profile millions of cells per tissue while preserving each cell’s spatial coordinates [14]. Two questions recur when representing these data: *what cell type or state is this cell?* and *what microenvironment is it sitting in?* A line of graph-based methods – NicheCompass [1], GraphST [12], BANKSY [18], CellCharter [23], Novae [2] – learns *continuous* spatial embeddings that recover cellular niches.

A complementary line explores *discrete* token vocabularies for tissue data, motivated by transformer foundation models that consume tokens rather than continuous vectors: cells are tokenised as gene-rank or binned-expression sequences [19, 24], as whole-cell projections [25], or via a fixed *k*-means “meta-cell” vocabulary [7]. VQ-VAE itself has been applied to non-spatial single-cell integration [11] and transcript point clouds in FISH imaging [28], and VQGraph [27] learns a structure-aware codebook on a GNN; however, none discretises the per-cell gene-expression matrix. Discrete tokens promise *interpretability* (human-readable identifiers tracked across samples), *stability* (no rotational ambiguity), *compactness* (a short tuple of integers per cell), and *compositionality* (token sequences consumable by downstream models).

The token-based and spatial-graph lines remain largely separate: token-based models either rely on heavy single-cell pretraining and fine-tune on SRT [19, 25, 24], or use a fixed *k*-means codebook blind to spatial context [7]. Any single-codebook design, the natural graph VQ-VAE [27], forces one codebook to represent both cellular identity and tissue context; learning two structurally disjoint codebooks is the natural fix and motivates SQUINT’s design.

### Contributions

We propose SQUINT, a graph vector-quantized variational autoencoder that learns separate cell and niche codebooks directly from spatial transcriptomics data. Concretely:

- **Dual discrete codebooks with residual VQ (Sections 3.2 and 3.3)**. An MLP trunk produces a per-cell latent quantised *directly* by a cell codebook **E**^cell^ (cell-intrinsic identity, before any neighbourhood mixing), while a GNN aggregates it into a neighbourhood latent quantised by a niche codebook **E**^niche^ (microenvironment). Each is a residual stack of *L* levels, giving a coarse-to-fine vocabulary learned directly from the spatial cell-by-gene matrix; the codes are compact, trackable cell- and niche-identifiers.
- **Domain-relevant design choices (Section 3.4**). A within-section cosine adjacency BCE anchors the niche codes in the spatial graph, a cross-section mutual-nearest-neighbour (MNN) contrastive loss on the cell latents aligns transcriptomically matched cells across sections, and a decoder section covariate absorbs batch effects; ablations show all three are essential (Section 4.6).
- **Empirical evaluation (Section 4)**. Across three datasets and four assays we benchmark SQUINT against spatial methods, single-cell foundation models and scVI, covering niche and cell-type identification, cross-section integration, held-out-region gene imputation, and one-step query-to-reference mapping on a human NSCLC cohort.

## 2. Related Work

### Continuous and discrete representations for SRT

Graph autoencoders learn *continuous* per-cell embeddings on a spatial *k*-NN or Delaunay graph and cluster them post-hoc (STAGATE [6], SpaceFlow [15], GraphST [12], SEDR [26]); NicheCompass [1] is the closest precedent to our spatial-prior choices (conditional VGAE, NB likelihood with library-size scaling, cosine-similarity adjacency BCE). *Discrete* cellular vocabularies have so far come either from transformer pretraining on scRNA-seq with gene-rank or binned-expression tokens (Nicheformer [19], scGPT-spatial [24]) or from fixed clustering of projected expression (GeST [7]).

### VQ-VAE and residual quantization

Learned VQ codebooks [21] are only beginning to reach single-cell data and, to our knowledge, have not been applied to the spatial cell-by-gene matrix: CVQVAE [11] quantises *non-spatial* multi-omics, [28] applies a VQ-VAE to FISH transcript and centroid coordinates rather than the count matrix, and learned tokenizers for *dissociated* scRNA-seq are a concurrent area. SQUINT instead operates on the same cell-by-gene matrix as the cell-type and niche baselines and is, to our knowledge, the first to learn *dual* cell/niche codebooks on it: our niche branch follows graph vector quantization (VQGraph [27], a VQ codebook on a GNN with EMA updates), which targeted GNN-to-MLP distillation rather than biological tokenisation, and a single-codebook port would inherit the cell-vs-niche conflation we avoid; residual vector quantization (RVQ) [29, 5] stacks *L* codebooks so that level *ℓ*+1 quantises the residual of levels 1:*ℓ*, a coarse-to-fine hierarchy without an exponentially large flat codebook.

## 3. Methods

### 3.1. Problem Setting and Notation

A spatial transcriptomics sample is a triple (**X**, *P*, **C**) where 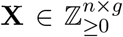 is the cell-by-gene count matrix, *P* ∈ *R*^*n*×2^ stacks the 2D spatial coordinates of the *n* cells, and **C** ∈ {0, 1}^*n*×*m*^ is the per-cell one-hot condition matrix, which in our experiments encodes the section identifier over *m* sections. We build a symmetrised *k*-nearest-neighbour spatial graph *G* = (*V, E*) over *P* (*V* = [*n*], adjacency **A** ∈ {0, 1}^*n*×*n*^). The goal is to learn, for each cell *c* ∈ *V*, a pair of discrete codes 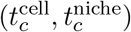 such that 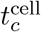 identifies cell-intrinsic identity (cell type or state) and 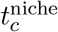 identifies the cell’s niche.

### 3.2. Dual Codebook Architecture

SQUINT consists of an MLP trunk, a graph encoder, two vector-quantization layers, and two negative-binomial decoders (Fig. 1). The forward computation is:

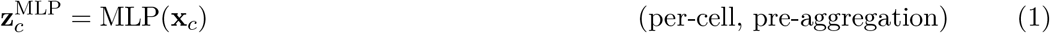

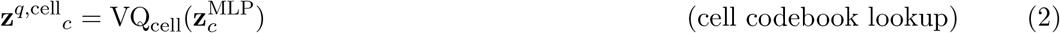

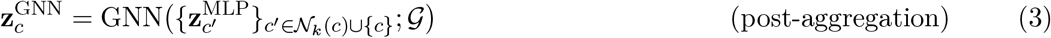

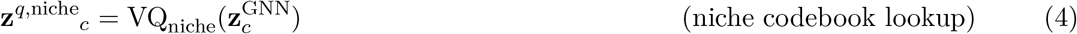

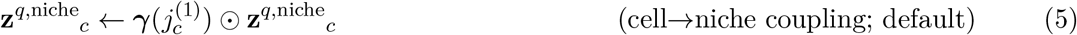

**Fig. 1:**
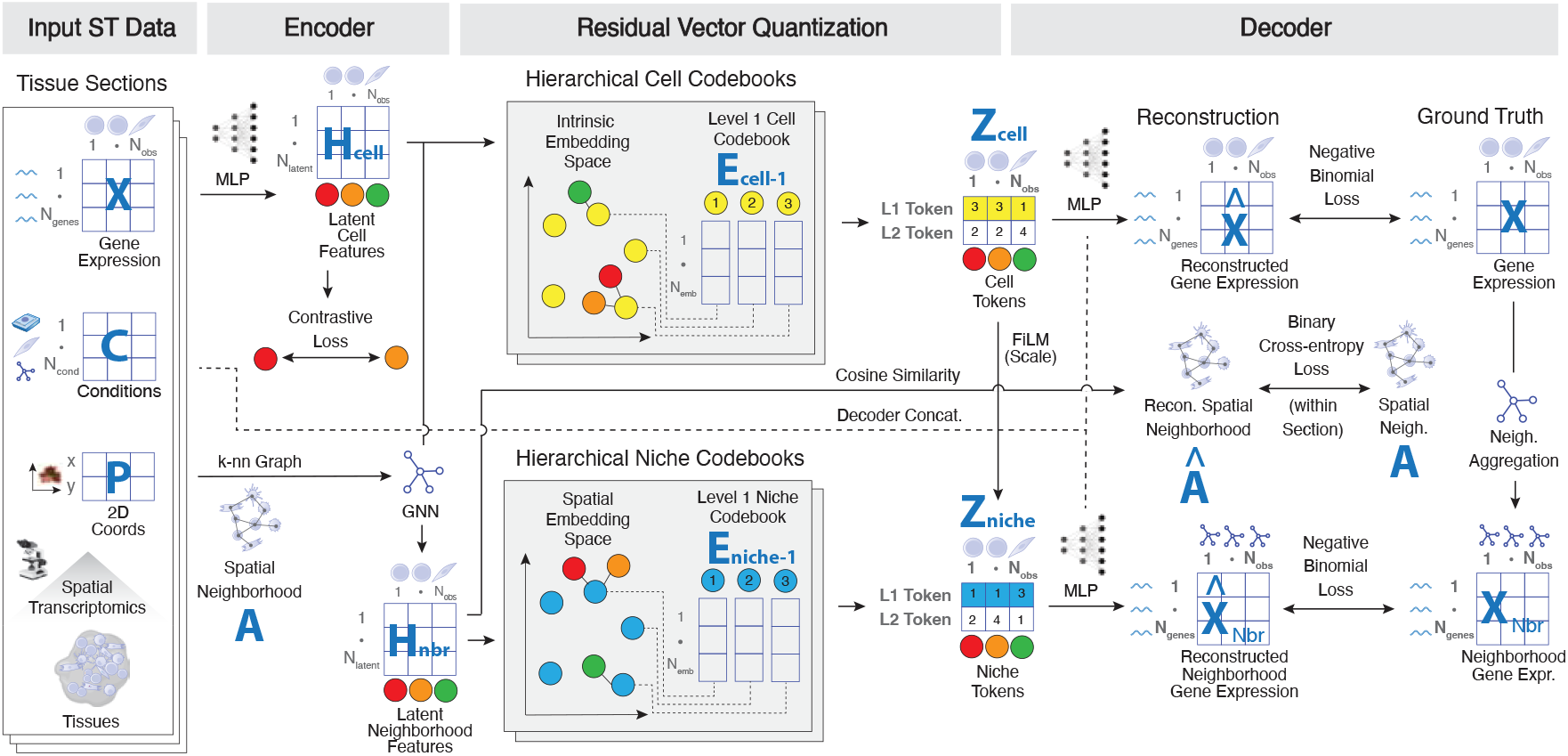
SQUINT architecture. An MLP encodes per-cell counts **X** into cell latents **H**^cell^; a GNN over the spatial *k*-NN graph yields neighbourhood latents **H**^nbr^. Two residual codebooks (**E**^cell^, **E**^niche^) quantise the branches and two negative-binomial decoders reconstruct per-cell and 1-hop-neighbourhood counts. Auxiliary objectives and the decoder section covariate **C** are detailed in Section 3; downstream tasks in Fig. S1.

Stacking the pre-aggregation latents 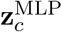 gives the *Latent Cell Features* **H**^cell^ ∈ R^*n*×*d*^ and stacking the post-GNN latents 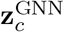 the *Latent Neighborhood Features* **H**^nbr^. Because the cell codebook quantises the MLP output *before* any aggregation, VQ_cell_ has no access to neighbours and is biased toward cell-intrinsic signal; because the niche codebook quantises the post-GNN output, VQ_niche_ sees neighbour-mixed signal and is biased toward spatial context. The last line (Equation (5)) is SQUINT’s default *cell* → *niche coupling*: 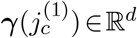 is a learned scale vector (initialised to **1**, scale-only) indexed by the cell’s level-1 code 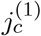, so the discrete cell type re-weights the niche dimensions before decoding; ***γ*** ≡ **1** recovers the decoupled variant (Table 3).

Two negative-binomial decoders reconstruct counts from the quantised codes: the cell decoder maps 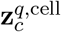 to raw per-cell counts 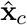, and the niche decoder maps 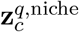 to a *per-cell* profile that is spatially averaged over *c*’s 1-hop neighbours to form the neighbourhood-mean prediction 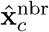, fit to the observed 1-hop mean; the decoder does not emit a neighbourhood profile directly (Appendix A). Both are scaled by the cell’s library size *ℓ*_*c*_ = ∑ _*j*_ *x*_*c,j*_ and conditioned on the section covariate **c**_*c*_ (Section 3.4), and each branch carries an *independent* per-gene inverse dispersion – a shared dispersion is dragged toward the smoother niche target and degrades the cell-branch likelihood. The decoder equations and the dispersion-decoupling rationale are in Appendix A.

### 3.3. Vector Quantization and the Residual Extension

Each branch quantises its embedding **z** ∈ R^*d*^ against a codebook **E** ∈ R^*K*×*d*^ by maximum-cosine-similarity lookup [21], trained with straight-through gradients and exponential-moving-average codebook updates (decay *β*=0.8) with dead-code resampling for collapse robustness [9]; codebook health is stable across *β* ∈ [0.5, 0.95] (Fig. S4c). We extend each codebook into a residual stack of *L* levels (RVQ) [29, 5]: level *ℓ*+1 quantises the residual left by levels 1:*ℓ*, so each cell receives a code tuple 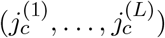 that forms a coarse-to-fine hierarchy. The default uses *L*=2 with codebook sizes (*K*^(1)^, *K*^(2)^) = (30, 90) per branch (justified in Table S1). The full VQ and RVQ equations, the cosine-VQ rationale, the stop-gradient detail, and the initialisation / dead-code-resampling scheme are given in Appendix A.

### 3.4. Domain-Relevant Objectives and Training

A vanilla VQ-VAE would quantise expression in isolation; SQUINT adds three lightweight, domain-motivated objectives – a spatial-structure loss, a cross-section correspondence loss, and a batch covariate – that write tissue geometry, cross-section invariance and batch awareness directly into the discrete vocabulary. Ablations show all three are essential (Section 4.6).

#### Within-section adjacency loss (spatial structure)

To anchor the niche codebook in the spatial graph, a cosine-similarity adjacency reconstruction loss *ℓ*_adj_ [1] applies a binary cross-entropy to the temperature-scaled cosine similarity of the *continuous* post-GNN embeddings **z**^GNN^ of balanced true-edge / non-edge cell pairs sampled within each section.

#### Cross-section contrastive loss (cross-section correspondence)

The cell-branch term *ℓ*_contr_ on the post-MLP cell latent **z**^MLP^ combines a within-section InfoNCE NT-Xent [22] (*k*_pos_=5 positive neighbours, in-batch negatives, temperature 0.1), which sharpens cell-type resolution, with a cross-section mutual-nearest-neighbour *pure-attraction* term (*k*_cross_=1, no negatives) that pulls each cell toward its mutual nearest neighbour in the *other* section(s), aligning transcriptomically matched cells across sections.

#### Decoder section covariate (batch awareness)

Concatenating the one-hot section identifier **c**_*c*_ with **z**^*q*,cell^ and **z**^*q*,niche^ before each decoder lets the decoders absorb per-section gene patterns, freeing the codebooks to encode section-invariant biology, the dominant batch-correction lever in our ablations (Table 3).

#### Optimization objective

SQUINT is trained end-to-end by minimizing

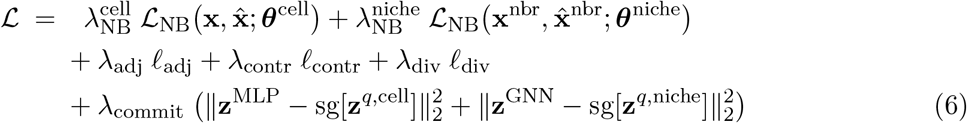

which sums the two per-branch NB likelihoods, the within-section adjacency BCE, the cross-section contrastive loss, a cell-branch codebook-diversity regulariser *ℓ*_div_, and the two VQ commitment terms; the codebooks are updated by EMA via the straight-through estimator. Loss weights, per-term expressions, and optimisation settings are in Appendices A and B.

### 3.5. Spatial Imputation via a Masked-Code Prior

Because a trained SQUINT maps every cell to a short stack of discrete tokens, gene-expression imputation in unmeasured regions reduces to predicting the *tokens* of held-out cells from the tokens of their observed spatial neighbours, and then decoding. We learn this with a lightweight second stage on top of the *frozen* stage-1 model: a graph-native, MaskGIT-style [3] masked-code transformer that operates purely on the discrete codes, reading the tokens of observed cells to predict those of the masked ones. Each cell is one ordered stack (the two cell and two niche RVQ levels); per-(branch, level) heads condition on shallower residual levels and, cross-branch, on the predicted cell stack, mirroring SQUINT’s cell → niche coupling [10]. Trained by masked cross-entropy on the frozen codes, the prior is made spatial rather than sequential by three adaptations – local-patch attention with a learnable distance bias, a random-Fourier coordinate encoding, and spatially-contiguous block masking (Appendix A.4). At inference we in-paint a held-out region by iterative, confidence-scheduled parallel decoding, obtaining a posterior over each held-out cell’s code stack. Rather than commit to a single path (the single decode, “SQUINT (No MC)”), “SQUINT (MC)” forms a *Monte-Carlo* estimate of E[**x** | context], the minimum-MSE predictor under the learned posterior: it draws *S*=1000 code configurations from these posteriors, decodes each through the frozen decoders, and averages the counts.

## 4. Experiments

We evaluate SQUINT on niche identification (from the niche codebook), cell-type identification (from the cell codebook), and cross-section data integration, including across assays (from the quantized cell and niche embeddings), plus gene-expression reconstruction/imputation and query-to-reference mapping (Fig. 3). Together these probe both halves of SQUINT’s dual-codebook contribution: whether the cell and niche codes recover the biology they were designed to capture, and whether the same model integrates cells across sections without per-batch retraining. The headline finding is that SQUINT *outperforms or is competitive with the leading continuous baselines* on every task, despite committing each cell to a small set of discrete tokens.

### 4.1. Setup

#### Datasets

We use one STARmap [17] and one MERFISH [31] mouse-brain section, each from a single donor (Fig. S1a), deliberately chosen to cover the *same anatomical region* so that cross-assay integration is well-posed: the sections share cell types and niches and *should* align, so residual separation reflects the assay (batch) effect rather than biology. Each ships with expert niche and cell-type annotations (Fig. S2a, b) against which NMI / ARI are computed per section and averaged. After standard preprocessing (cell-by-gene counts, *k*=16 spatial *k*-NN graph), SQUINT is trained jointly on both donors. To test generalisation, we also benchmark identification on two single-assay human datasets (Table 1): CosMx NSCLC (two sections, one donor; the full cohort is used for query-to-reference mapping in Section 4.5) and Xenium Eczema (skin; three sections, different donors), integrating across each dataset’s sections.

**Table 1:**
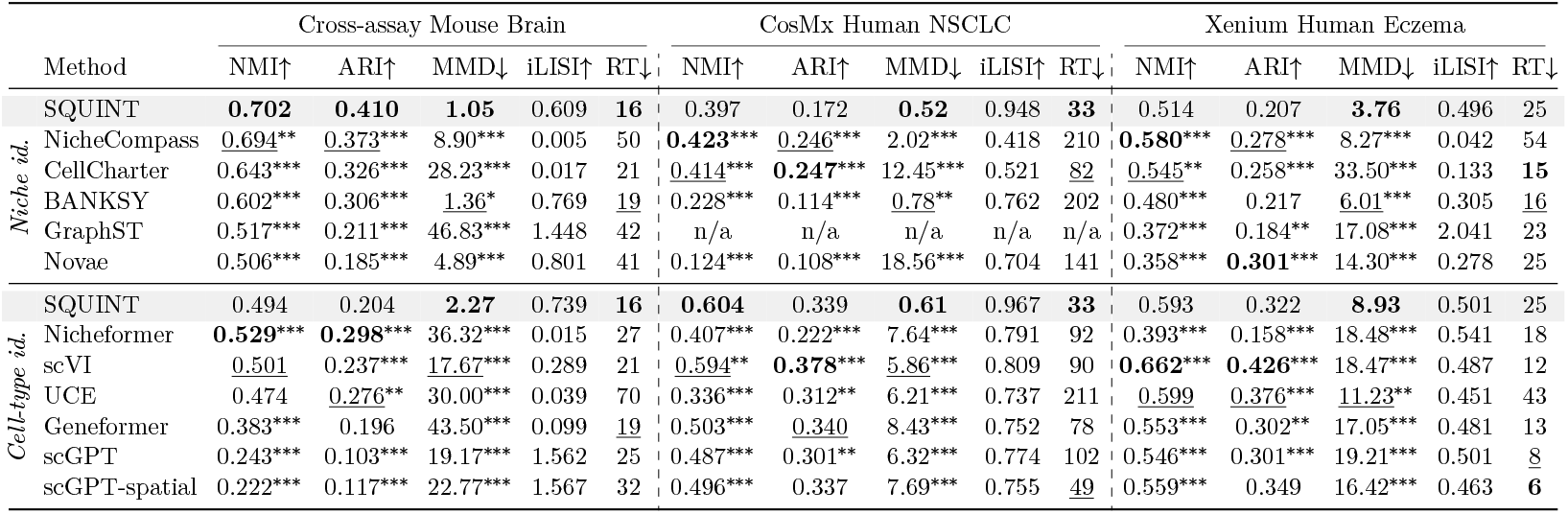
Niche- and cell-type-identification benchmark across three datasets. Clustering quality (NMI, ARI), integration (MMD in 10^−3^; iLISI) and runtime (RT, min) for SQUINT (shaded) versus spatial-niche (top) and single-cell (bottom) methods. Five-seed means; **best**/second per column for NMI/ARI/MMD/RT. Superscripts on the NMI/ARI/MMD entries give each method’s significance vs. the shaded SQUINT row (Welch’s *t*-test across seeds; *p<0.05, **p<0.01, ***p<0.001). `n/a`: GraphST out-of-memory on CosMx.

#### Baselines

For niche identification we compare against spatial-context methods (NicheCompass [1], CellCharter [23], BANKSY [18], GraphST [12], Novae [2]); for cell-type identification against the strong non-foundation baseline scVI [13] and the single-cell foundation models Nicheformer [19], UCE [16], Geneformer [20], scGPT [4] and scGPT-spatial [24]; integration is evaluated on SQUINT’s VQ-quantized cell and niche embeddings (**z**^*q*,cell^, **z**^*q*,niche^) and on each baseline’s latent embedding.

Per-method descriptions and the post-hoc Harmony [8] / preprocessing-time PASTE [30] alignment used by BANKSY and GraphST are in Appendix C.

#### Metrics

Identification quality is NMI and ARI against the expert annotation: SQUINT’s Level-1 token *K*^(1)^=30 *is* the label, whereas continuous baselines need a Leiden step to the same 30-cluster granularity, counted in their runtime – a fair, matched-granularity comparison. Cross-section integration uses Maximum Mean Discrepancy (MMD; ↓) and the integration-Local Inverse Simpson Index (iLISI; ↑), which together separate faithful integration from over-integration (high iLISI with high MMD). We also report runtime (RT) on a single NVIDIA H200 GPU. The clustering protocol and MMD / iLISI settings are in Appendix C.

### 4.2. Niche Identification and Cross-Section Integration

Table 1 (top) shows SQUINT leads niche NMI and ARI on the cross-assay Mouse Brain; on CosMx NSCLC and Xenium Eczema it trails the best clustering baselines on NMI/ARI but attains the lowest MMD (best cross-section integration) on every dataset at competitive runtime, despite only a few discrete niche tokens per cell.

On NSCLC it trains in 33 min, against 82–210 min for the two methods that edge it on NMI. Fig. 2 confirms this qualitatively: SQUINT’s Level-1 niche codes recapitulate the Allen Reference Atlas compartments and the ground-truth labels (Fig. S2a) across both assays, with no clustering. On the Mouse Brain – the only dataset pairing two assays – SQUINT also attains the lowest cross-assay MMD (1.05 × 10^−3^) without sacrificing niche clustering, the signature of *faithful* integration. A high iLISI alone does not establish this: GraphST’s far higher iL-ISI comes from PASTE [30] preprocessing but pairs with weak NMI/ARI/MMD – over-integration, not alignment (Fig. S3b).

**Fig. 2:**
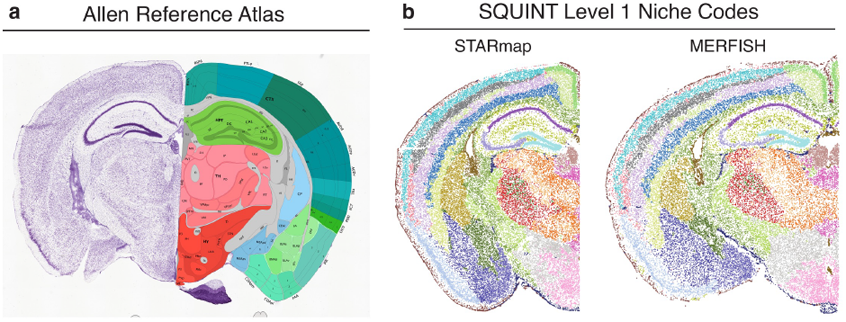
Brain anatomy **(a)** is recapitulated by SQUINT **(b)**.

**Fig. 3:**
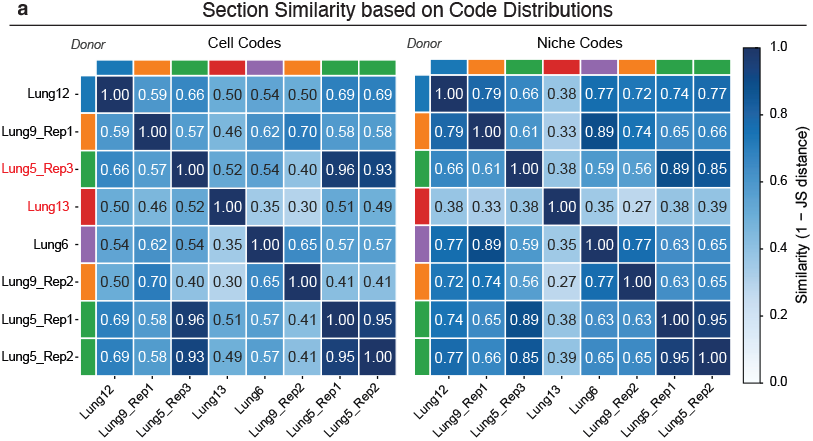
SQUINT query-to-reference mapping (CosMx NSCLC): cell- and niche-code distribution similarity (1 − JS); queries in red.

### 4.3. Cell-Type Identification and Cross-Section Integration

Table 1 (bottom) shows SQUINT is competitive on cell-type identification with scVI [13] and the large single-cell foundation models (Nicheformer, UCE, Geneformer, scGPT), which are pretrained on millions of cells: it attains the best cell-type NMI on CosMx NSCLC, comes within a few points of the best on the Mouse Brain (0.49 vs. 0.53), and trails scVI/UCE on Xenium (0.59 vs. 0.66/0.60). As on the niche branch, its advantage is integration and speed: the lowest cell-branch MMD on all three datasets and the fastest runtime on the Mouse Brain and NSCLC, whereas scGPT and scGPT-spatial reach high iLISI on the Mouse Brain but the weakest NMI/ARI and high MMD – the classic over-integration signature (Fig. S3d). SQUINT’s speed follows from its discrete tokens serving directly as cluster assignments (no Leiden step).

### 4.4. Gene-Expression Reconstruction and Spatial Imputation

SQUINT’s discrete tokens support gene-expression *reconstruction* (each held-out cell encoded from its *own* counts and rebuilt at the per-cell and 1-hop level) and true *imputation* (held-out cells reconstructed *without reading their expression*, their tokens predicted from observed neighbours by a separate second-stage masked-code prior over the frozen codes (Section 3.5; Fig. S1d, e). Both are scored by Pearson and Spearman correlation, RMSE and expressed-gene detection (AUROC, AP), over all genes and highly-variable subsets (Appendix C).

Reconstruction baselines are NicheCompass [1], scVI [13] (per-cell and neighbourhood-aggregated variants), and a single-codebook *Vanilla VQ-VAE* ablation; imputation compares SQUINT (MC/No MC) against GeST [7] and a non-parametric spatial-*k*NN floor (Appendix C).

#### Results

On reconstruction (Table 2, top) SQUINT leads most neighbourhood-level metrics and is competitive at the cell level – notable given its discrete 30 × 90-code bottleneck and 32-unit decoder (Appendix B). On held-out imputation (bottom), scored leak-free (each held-out cell’s library size comes from its *observed* neighbours, as for GeST/*k*NN; Appendix C), Monte-Carlo averaging beats the single decode on every metric, significantly on most, leading at both levels bar cell-wise RMSE (GeST’s smoother). Fig. S5 shows qualitative reconstructions.

**Table 2:**
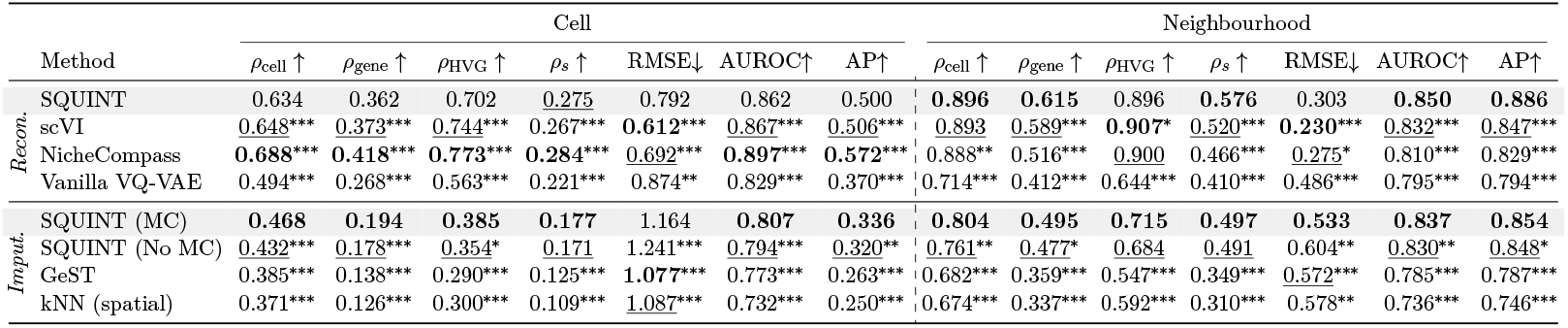
Gene-expression reconstruction and imputation at the cell (per-cell) and neighbourhood (1-hop) levels on mouse-brain STARmap + MERFISH. *Reconstruction*: held-out cells encoded from their own counts, SQUINT (shaded) versus continuous baselines. *Imputation*: held-out regions predicted without reading their expression, by SQUINT (MC), a second-stage masked-code prior over the frozen SQUINT codes with *S*=1000 Monte-Carlo decoding (shaded), versus the single-decode SQUINT (No MC), GeST and a spatial-*k*NN floor. Metrics: Pearson *ρ* (cell-, gene-wise, HVG) and Spearman *ρ*_*s*_ on log1p, RMSE on raw counts, expressed-gene detection (AUROC, AP). Five-seed means; **best**/second per column *within each block*. Superscripts give significance vs. the shaded reference (SQUINT for reconstruction, SQUINT (MC) for imputation; Welch’s *t*-test across seeds; ^*^*p<*0.05, ^**^*p<*0.01, ^***^*p<*0.001).

### 4.5. Query-to-Reference Mapping on CosMx Human NSCLC

A discrete representation makes query-to-reference (atlas) mapping natural: each section is summarised by the empirical distribution of its cell- and niche-tokens, so section similarity is a distance between two categorical distributions – no continuous re-embedding or retraining at query time.

On a CosMx non-small-cell lung cancer (NSCLC) cohort (five donors, eight samples), SQUINT is trained on a reference (four donors, six samples); the frozen model then assigns Level-1 codes to two held-out query samples, from a seen and an unseen donor. The seen-donor replicate (Lung5_Rep3) matches its donor’s other replicates (cell-code similarity 0.96/0.93), while the unseen donor (Lung13) has only weak matches (≤ 0.52), as expected. This matching is meaningful because each Level-1 code is a stable, interpretable identifier: it tracks a consistent expert cell type or niche across donors and replicates (Fig. S6a–d), so a section’s code histogram is a biological fingerprint rather than a batch-specific embedding, and same-donor sections share compositions (Fig. S6e, f).

### 4.6. Ablations

We ablate the three recipe-specific components on the STARmap + MERFISH benchmark (Table 3). Removing the adjacency loss collapses niche NMI / ARI and weakens niche iLISI / MMD, confirming it anchors the niche codebook. Replacing the cross-section MNN contrastive with a within-section variant collapses cell-branch integration (cell iLISI 0.74 → 0.49), whereas removing it entirely preserves integration but degrades cell-type identification (cell NMI 0.49 → 0.42). Removing the decoder section-ID covariate inflates cell MMD by an order of magnitude and collapses cell iLISI, confirming it is the primary batch-correction lever. Table 3 further isolates three axes. Clustering the latent instead of using the codes – the VQ model’s own (VQ (Leiden)) or a continuous VAE’s (VAE (Leiden)) – can edge the codes on NMI (Leiden optimises the partition directly) yet collapses integration (niche MMD 1.05 → 16–20 × 10^−3^, iLISI 0.61 → 0.04–0.05): even the same VQ latent breaks once clustered, so discreteness, not architecture, drives faithful integration. Flattening the two-level (30, 90) stack to one codebook of equal capacity (*K*=2700) preserves niche identification but guts the cell branch (cell ARI 0.20 → 0.01): coarse-to-fine factorisation, not raw size, drives cell quality. Across coupling variants NMI/ARI are near-tied, but FiLM scaling gives the strongest niche mixing (iLISI 0.61 vs. ≤ 0.36) at comparable MMD; Table S1 sweeps capacity.

**Table 3:**
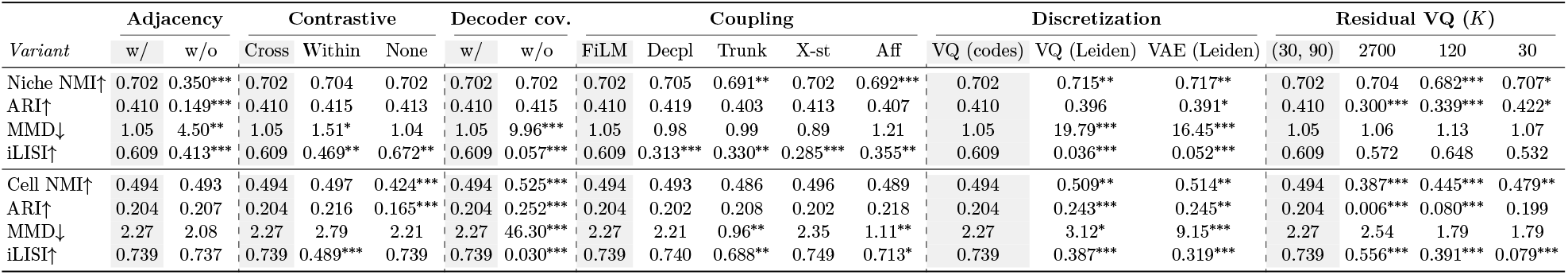
Ablations. Training-recipe and bottleneck axes on mouse-brain STARmap + MERFISH; each column is a variant grouped by axis, shaded = default. Five-seed means; superscripts give significance vs. default (^*^/^**^/^***^); MMD in 10^−3^. Coupling: FiLM(=cell-cond. scale), Decpl, Trunk, X-st(=cross-stitch), Aff; Discretization: VQ (codes) uses the codes directly; VQ (Leiden)/VAE (Leiden) Leiden-cluster the VQ or continuous-VAE latent to *K*=30. Leiden optimises the partition directly, so it can edge the codes’ NMI at matched *K*; the codes’ advantage is integration, not clustering quality. Residual VQ (*K*): (30, 90) default; flat codebooks of matched *product* (2700=30 *×* 90) or *sum* (120=30+90) capacity; one level (30). Capacity sweeps in Table S1.

## 5. Conclusion

We presented SQUINT, a graph VQ-VAE that learns disjoint cell and niche residual codebooks from spatial transcriptomics. Across three datasets spanning four assays (STARmap, MERFISH, CosMx, Xenium), SQUINT is competitive with strong baselines on niche and cell-type identification and achieves the most faithful cross-section integration (lowest MMD everywhere) at low cost. On the cross-assay mouse brain it further imputes held-out regions, and on a CosMx Human NSCLC cohort the same tokens enable one-step query-to-reference mapping. Ablations confirm every component matters, the domain-relevant losses especially; the discrete codes form a compact, interpretable token vocabulary for downstream transformer pretraining.

