## Supplementary Material for "Learning Discrete Cell and Niche Codes from Spatial Transcriptomics Using Dual Residual Vector Quantization"

#### Appendix A. Extended Methods

##### A.1. Vector Quantization and the Residual Extension

For an embedding $\mathbf{z} \in \mathbb{R}^d$ and a codebook $\mathbf{E} \in \mathbb{R}^{K \times d}$, the standard VQ-VAE operation [18] is

$$j^\star = \arg\max_{j \in [K]} \cos(\mathbf{z}, \mathbf{e}_j), \qquad \mathrm{VQ}(\mathbf{z}) = \mathbf{e}_{j^\star}, \tag{A.1}$$

where we use cosine similarity rather than Euclidean distance for improved codebook usage and collapse robustness [22], and the gradient is propagated back to $\mathbf{z}$ via the straight-through estimator. The codebook itself is updated by an EMA of assigned encoder outputs (decay $\beta = 0.8$, deliberately faster than the 0.99–0.999 typical of large codebooks: our small 30/90-code levels receive many assignments per code each step, so a shorter averaging window tracks the moving encoder distribution and stabilises the dead-code rate) rather than by gradient. Codes whose EMA cluster mass falls below a threshold $\tau$=2 are re-initialised from current encoder outputs to avoid codebook collapse [9]. Codebooks are initialised by $k$-means on a sample of encoder outputs from the first batch. The resulting per-level code-usage distributions and perplexities on the mouse-brain benchmark are reported in Fig. S4.

**Residual VQ.** Either codebook can be extended into a residual stack of $L$ codebooks $\{\mathbf{E}^{(\ell)}\}_{\ell=1}^{L}$ of sizes $K^{(\ell)}$, following neural audio codecs [23, 5]. Iteratively:

$$\mathbf{c}^{(\ell)} = \mathrm{VQ}^{(\ell)}(\mathbf{r}^{(\ell)}), \qquad \mathbf{r}^{(\ell+1)} = \mathbf{r}^{(\ell)} - \mathrm{sg}[\mathbf{c}^{(\ell)}], \qquad \mathbf{r}^{(1)} = \mathbf{z}, \tag{A.2}$$

with the final quantised vector $\mathbf{z}^q = \sum_{\ell=1}^{L} \mathbf{c}^{(\ell)}$ and the per-cell discrete code being the tuple $(j_c^{(1)}, \ldots, j_c^{(L)})$. The stop-gradient on the residual update is essential: without it, the straight-through estimator at level 1 zeroes the gradient at all deeper levels [23]. We enable $k$-means initialisation and dead-code resampling only on level 1; deeper levels see residuals that are not directly clusterable from the data distribution. The default configuration uses $L$=2 levels per branch with codebook sizes $(K^{(1)}, K^{(2)}) = (30, 90)$ on both the cell and niche branches (justified in Table S1); smaller $L$ collapses the hierarchy to a single codebook and larger $L$ gives an even finer coarse-to-fine token tree.

##### A.2. Negative-Binomial Decoders

The cell decoder reconstructs raw per-cell counts:

$$\hat{\mathbf{x}}_c = \mathrm{Dec}_{\mathrm{cell}}(\mathbf{z}_c^{q,\mathrm{cell}}, \mathbf{c}_c) \cdot \ell_c, \qquad \mathbf{x}_c \mid \mathbf{z}_c^{q,\mathrm{cell}} \sim \mathrm{NB}(\hat{\mathbf{x}}_c, \boldsymbol{\theta}^{\mathrm{cell}}), \tag{A.3}$$

where $\ell_c = \sum_{j=1}^{g} x_{c,j}$ is the cell's library size, $\mathbf{c}_c$ the per-cell section covariate (Section 3.4), and $\boldsymbol{\theta}^{\mathrm{cell}} \in \mathbb{R}^g_{>0}$ a per-gene inverse dispersion. The niche decoder uses the same $\ell_c$ and reconstructs the $r$-hop neighbourhood-aggregated counts:

$$\tilde{\mathbf{x}}_c = \mathrm{Dec}_{\mathrm{niche}}(\mathbf{z}_c^{q,\mathrm{niche}}, \mathbf{c}_c) \cdot \ell_c, \qquad \mathbf{x}_c^{\mathrm{nbr}} \mid \mathbf{z}_c^{q,\mathrm{niche}} \sim \mathrm{NB}(\hat{\mathbf{x}}_c^{\mathrm{nbr}} = \mathrm{Agg}_r(\tilde{\mathbf{x}})_c, \boldsymbol{\theta}^{\mathrm{niche}}), \tag{A.4}$$

so the niche decoder emits a *per-cell* profile $\tilde{\mathbf{x}}_c$, *not* a neighbourhood profile; the neighbourhood-mean prediction $\hat{\mathbf{x}}_c^{\mathrm{nbr}}$ is obtained by aggregating these per-cell outputs. Here $\mathrm{Agg}_r$ is the $r$-hop spatial mean over the true, fixed $k$-NN graph $\mathcal{G}$ ($r$=1): each cell's per-cell prediction $\tilde{\mathbf{x}}_c$ is averaged over its true spatial neighbours and compared to the mean of the *observed* counts over the same neighbours; the aggregation never uses a reconstructed neighbourhood, and $\mathcal{G}$ is built once from $\mathcal{P}$ and held fixed. Decoupling the dispersions is empirical: the per-cell and neighbourhood-mean counts have very different variance structure, and a single shared $\boldsymbol{\theta}$ is dragged toward the smoother niche target, degrading the cell-branch likelihood throughout training.

##### A.3. Auxiliary Objectives

**Within-section adjacency loss (Section 3.4).** For positive (true-edge) and negative (non-edge) cell pairs sampled from the intra-section graph,

$$\ell_{\mathrm{adj}} = \mathrm{BCE}\Big(\tfrac{1}{\tau}\cos(\mathbf{z}_i^{\mathrm{GNN}},\, \mathbf{z}_j^{\mathrm{GNN}}),\;\; \mathbf{1}[(i,j)\in\mathcal{E}]\Big), \tag{A.5}$$

with cosine temperature $\tau = 0.1$ so the logits span roughly $[-10, +10]$ and the BCE can saturate near 0/1. Using the continuous $\mathbf{z}^{\mathrm{GNN}}$ rather than the quantised $\mathbf{z}^{q,\mathrm{niche}}$ keeps a gradient between cells that share a niche code (whose codes have cosine similarity exactly 1) and matches NicheCompass, inheriting spatial structure into the codebook when $\mathbf{z}^{\mathrm{GNN}}$ is discretised by $\mathrm{VQ}_{\mathrm{niche}}$.

**Cell-branch contrastive loss (Section 3.4).** The cell-branch term combines a within-section InfoNCE NT-Xent and a cross-section mutual-nearest-neighbour attraction term on $\mathbf{z}^{\mathrm{MLP}}$: $\ell_{\mathrm{contr}} = \ell_{\mathrm{NT}} + \lambda_{\mathrm{cross}}\,\ell_{\mathrm{MNN}}$. With within-section $k_{\mathrm{pos}}$-nearest-neighbour positive set $\mathcal{P}_c$ and mini-batch $\mathcal{B}$ (all other batch cells act as negatives), the NT-Xent term [19] is

$$\ell_{\mathrm{NT}} = -\frac{1}{|\mathcal{B}|}\sum_{c\in\mathcal{B}}\frac{1}{|\mathcal{P}_c|}\sum_{p\in\mathcal{P}_c}\log\frac{\exp\big(\cos(\mathbf{z}_c^{\mathrm{MLP}},\mathbf{z}_p^{\mathrm{MLP}})/\tau_{\mathrm{c}}\big)}{\sum_{n\in\mathcal{B}\setminus\{c\}}\exp\big(\cos(\mathbf{z}_c^{\mathrm{MLP}},\mathbf{z}_n^{\mathrm{MLP}})/\tau_{\mathrm{c}}\big)}, \tag{A.6}$$

and, with $\mathcal{M}_c$ the cross-section mutual-nearest-neighbour set of anchor $c$, the pure-attraction term (no negatives) is

$$\ell_{\mathrm{MNN}} = -\frac{1}{|\mathcal{B}|}\sum_{c\in\mathcal{B}}\frac{1}{|\mathcal{M}_c|}\sum_{m\in\mathcal{M}_c}\cos\big(\mathbf{z}_c^{\mathrm{MLP}},\mathbf{z}_m^{\mathrm{MLP}}\big). \tag{A.7}$$

We use $k_{\mathrm{pos}} = 5$, $k_{\mathrm{cross}} = 1$, $\tau_{\mathrm{c}} = 0.1$, and $\lambda_{\mathrm{cross}} = 10$.

##### A.4. Masked-Code Prior (Stage 2)

The stage-2 prior (Section 3.5) is a graph-native MaskGIT-style [3] transformer over the frozen stage-1 codes. Three adaptations make it spatial rather than sequential: full self-attention within a local spatial patch with an additive, learnable per-head distance bias; a 2D random-Fourier encoding of (patch-normalised) coordinates; and spatially-*contiguous* block masking that matches the held-out-region geometry. Per-(branch, level) heads condition on shallower residual levels and, cross-branch, on the predicted cell stack, following stage-2 priors over residual codes [10]. At inference we in-paint a held-out region with MaskGIT's iterative, confidence-scheduled parallel decoding (cosine schedule, 20 steps, sampling temperature 0.5, no confidence noise), yielding a posterior over each held-out cell's code stack. The Monte-Carlo estimate ("SQUINT (MC)") draws $S$=1000 code configurations from these posteriors, decodes each through the frozen stage-1 decoders, and averages the resulting count profiles – an estimate of $\mathbb{E}[\mathbf{x} \mid \text{context}]$, the minimum-MSE predictor under the learned posterior; the single-decode ablation ($S$=1) is "SQUINT (No MC)".

#### Appendix B. Implementation Details

**Architecture.** The per-cell encoder is a three-layer MLP (hidden widths 400/400/256); the niche branch adds a single GNN layer (width 256) over the spatial $k$-NN graph. Each branch quantises a 256-dimensional embedding with a two-level residual codebook $(K^{(1)}, K^{(2)})=(30, 90)$. The two negative-binomial decoders (Equations (A.3) and (A.4)) are intentionally small — each a single 32-unit hidden layer mapping the 256-dimensional quantised code to the $g$ gene outputs — so that reconstruction quality reflects the discrete bottleneck rather than decoder capacity.

**Loss weighting.** In the training objective (Equation (6) of the main text) the per-branch NB log-likelihoods use the adjacency BCE of Equation (A.5) and the cell-branch contrastive loss of Equation (A.6); the two VQ commitment terms enter via the straight-through estimator, as the codebooks are updated by EMA rather than gradient. The cosine BCE has small per-pair magnitude ($\sim$ 0.5–1 nat at initialisation, bounded by $\log 2$) and would otherwise be swamped by the much larger gene-NB terms ($\sim$ 150 nats per cell); we up-weight it with $\lambda_{\text{adj}} = 1000$ so the spatial-structure signal shapes the niche codes materially rather than being dominated, and set $\lambda_{\text{contr}} = 10$. On the cell branch we add the codebook-diversity regulariser $\ell_{\text{div}}$ — the softmax-similarity code-repulsion term (temperature 100) of the main-text objective, Equation (6) — at the same weight, $\lambda_{\text{div}} = 10$, to encourage uniform codebook usage.

**Optimisation and mini-batch training.** We train with a seed-batch of $|\mathcal{B}| = 512$ cells per step on a spatial $k$-NN graph with $k = 16$ neighbours, using the `NeighborLoader` of PyTorch Geometric [6] configured with `num_neighbors=[16]` for the single GNN layer, i.e. no sub-sampling: each seed cell's full one-hop neighbourhood is loaded into the induced sub-graph for both message passing and adjacency BCE. The model is optimised with Adam (learning rate $7 \times 10^{-4}$, weight decay $10^{-3}$) for up to 80 epochs.

**Code Availability.** The model code is released as the `squint` Python package at https://github.com/Lotfollahi-lab/squint. The code that reproduces the experiments in this paper – benchmarking against all baselines, the ablations of Tables 3 and S1, and the CosMx Human NSCLC query-to-reference analysis – is provided in a separate repository at https://github.com/Lotfollahi-lab/squint-reproducibility, with configuration files for each variant reported in the manuscript.

**Data Availability.** All datasets used in this paper are publicly available. The mouse-brain MERFISH section [25] is distributed via CELLxGENE at https://cellxgene.cziscience.com/collections/0cca8620-8dee-45d0-aef5-23f032a5cf09. The mouse-brain STARmap section [14] is available on Zenodo at https://zenodo.org/records/8327576. The CosMx human non-small-cell lung cancer (NSCLC) FFPE dataset used in the query-to-reference mapping analysis is released by Bruker Spatial Biology at https://brukerspatialbiology.com/products/cosmx-spatial-molecular-imager/ffpe-dataset/nsclc-ffpe-dataset/.

**Funding.** S. Birk is supported by the Helmholtz Association under the joint research school MUDS.

**Generative AI Disclosure.** The authors used Anthropic's Claude (Opus 4.7) as a writing assistant during drafting and revision of this manuscript, including for prose, figure captions, and LaTeX formatting. All scientific content – claims, results, citations, and final phrasing – was reviewed, edited, and approved by the authors, who take full responsibility for the work. Generative AI was not used to generate references or to produce substantive unreviewed text.

#### Appendix C. Experimental Details

**Baselines.** For niche identification we use methods designed around spatial context: NicheCompass [1] (continuous VGAE with the same NB likelihood as SQUINT), CellCharter [20] (probabilistic clustering of GMM latents), BANKSY [15] (non-deep neighbourhood-augmented kernel clustering), GraphST [11] (graph-contrastive), and Novae [2] (graph-foundation-model). For cell-type identification we use scVI [12] – the variational autoencoder originally introduced as a deep generative model for single-cell RNA-seq, here run with a negative-binomial likelihood and included as a strong non-foundation baseline – alongside large single-cell foundation models pretrained on millions of cells: Nicheformer [16] (single-cell + spatial transformer foundation model), UCE [13] (Universal Cell Embeddings), Geneformer [17], scGPT [4], and the spatial variant scGPT-spatial [21]. For data integration we reuse the same baseline sets. BANKSY and GraphST do not natively integrate across batches: BANKSY relies on Harmony [8] *post-hoc* on the learned embedding, while GraphST runs on top of PASTE [24] coordinate alignment of the two sections *as a preprocessing step before training.* For the spatial gene-imputation task (Section 4.4) the baselines are NicheCompass [1] (continuous graph-VAE that, like SQUINT, reconstructs both per-cell and neighbourhood-aggregated counts), scVI [12] (single-cell NB likelihood without spatial information; we report both its native per-cell prediction and an "$\mathbf{X}_{\text{nbr}}$" variant trained directly on the 1-hop neighbourhood-aggregated counts so its output represents the local microenvironment), and a *Vanilla VQ-VAE*, the single-codebook ablation of SQUINT that removes the dual cell/niche split and uses one shared codebook for both reconstruction heads. For held-out-region *imputation* we additionally compare against GeST [7] (a generative model of spatial cellular context) and a non-parametric spatial-$k$NN floor, which predicts each held-out cell as the distance-weighted mean ($k$=16) of its nearest observed (train) cells' raw counts within the same section – a leave-one-out spatial smoother with no learned parameters.

**Metrics.** Niche- and cell-type-identification quality are measured by Normalised Mutual Information (NMI) and Adjusted Rand Index (ARI) between the predicted discrete labels and the ground-truth annotation. SQUINT provides discrete tokens directly: the per-cell prediction is the Level-1 RVQ token (the $K^{(1)} = 30$ coarse macro-cluster), used as the cluster assignment with no further clustering. For baselines that produce continuous embeddings we run Leiden clustering, sweeping the resolution to recover the same number of clusters as SQUINT's Level-1 codebook ($K^{(1)} = 30$), so that all methods are compared at matched cluster granularity. Cross-section data-integration quality is measured by the Maximum Mean Discrepancy (MMD; lower is better), computed separately on the cell and the niche embeddings. To make MMD comparable across embeddings of different dimensionality and scale, every embedding is first standardised (zero mean, unit variance per dimension) and the RBF kernel bandwidth is set per embedding by the median heuristic (the median pairwise distance on a subsample), so the reported MMD reflects distributional misalignment rather than raw coordinate scale. The complementary integration-Local Inverse Simpson Index (iLISI; higher is better) is reported in Fig. S3; the MMD / iLISI pair distinguishes faithful integration from over-integration – a method can inflate iLISI by mixing batches at the cost of biological signal, while MMD penalises distributional misalignment regardless of clustering.

**Reconstruction protocol and metrics (Section 4.4).** Held-out cells are tagged as a test split and never used as reconstruction targets or as seed cells, so neither decoder is fit on them; the spatial $k$-NN graph is built once over *all* cells and held fixed, and the identical train/test mask is applied to SQUINT and every baseline (a shared holdout routine), making the comparison like-for-like. At inference the frozen model encodes each held-out cell from its own counts – with per-cell library size computed from those counts, so no statistic is shared across cells – and the two assays interact only through the shared weights, not through any direct information transfer. Quality uses two Pearson families: the *gene-wise* correlation (per gene across cells, then averaged) measures recovery of each gene's spatial pattern, and the *cell-wise* correlation (per cell across genes, then averaged) measures recovery of each cell's full profile. Each is reported at the *cell level* (predicted per-cell counts $\hat{\mathbf{x}}_c$ vs. true $\mathbf{x}_c$) and the *neighbourhood level* (neighbourhood-mean prediction $\hat{\mathbf{x}}_c^{\text{nbr}}$ vs. true 1-hop mean $\mathbf{x}_c^{\text{nbr}}$).

For held-out-region *imputation*, where a cell's own counts are unavailable, we score every method *leak-free*: the decoder's library-size factor $\ell_c$ for a held-out cell is set to the mean library size of its 16 nearest *observed* (train) cells in the same section – never its own total counts – and neighbourhood-level scores aggregate each method's per-cell prediction over the spatial graph. Thus neither the target cell's sequencing depth nor its expression enters the prediction, exactly as for GeST and the $k$NN floor. (This matters for raw-count RMSE and expressed-gene detection, which are sensitive to library-size scaling; the correlation metrics are largely scale-invariant.)

### Appendix D. Additional Results

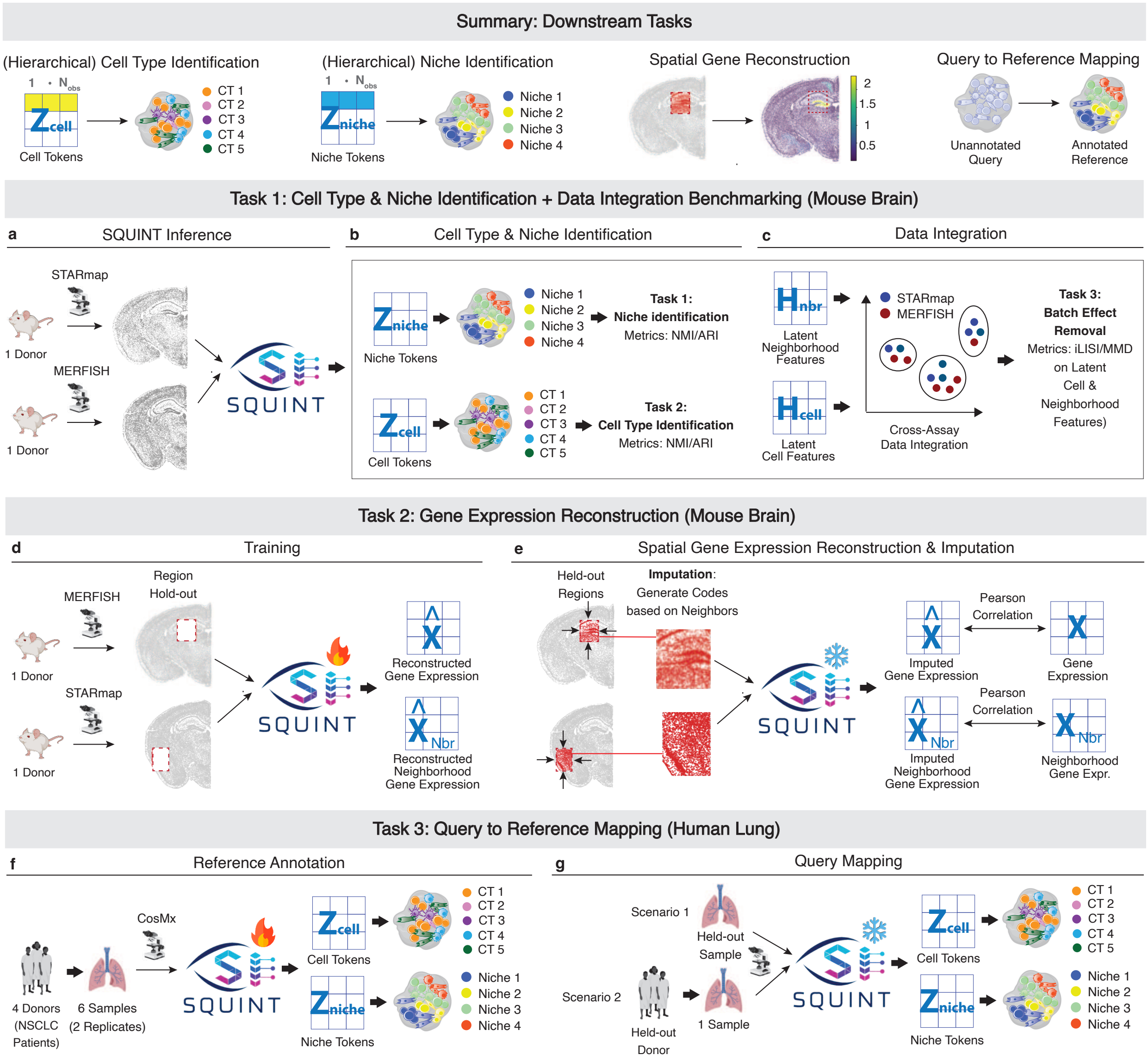


Fig. S1: Overview of the downstream tasks and their data. **(Top)** The discrete SQUINT codes support hierarchical cell-type and niche identification (from the cell and niche tokens), spatial gene-expression imputation, and query-to-reference mapping of an unannotated query onto an annotated reference. **Task 1 (mouse brain, a–c):** trained jointly on one STARmap and one MERFISH section (a); cell and niche tokens give cell-type and niche identification (NMI / ARI) (b) and cross-assay integration (iLISI, MMD) (c). **Task 2 (mouse brain, d–e):** trained with contiguous regions held out of both reconstruction targets (d); the frozen model reconstructs the held-out cells, scored at the cell and 1-hop neighbourhood level by Pearson correlation (e). **Task 3 (human lung, f–g):** trained on a CosMx Human NSCLC reference (four donors, six samples) (f); the frozen model maps held-out queries, a seen-donor replicate and an unseen donor, by comparing code-distributions (g).

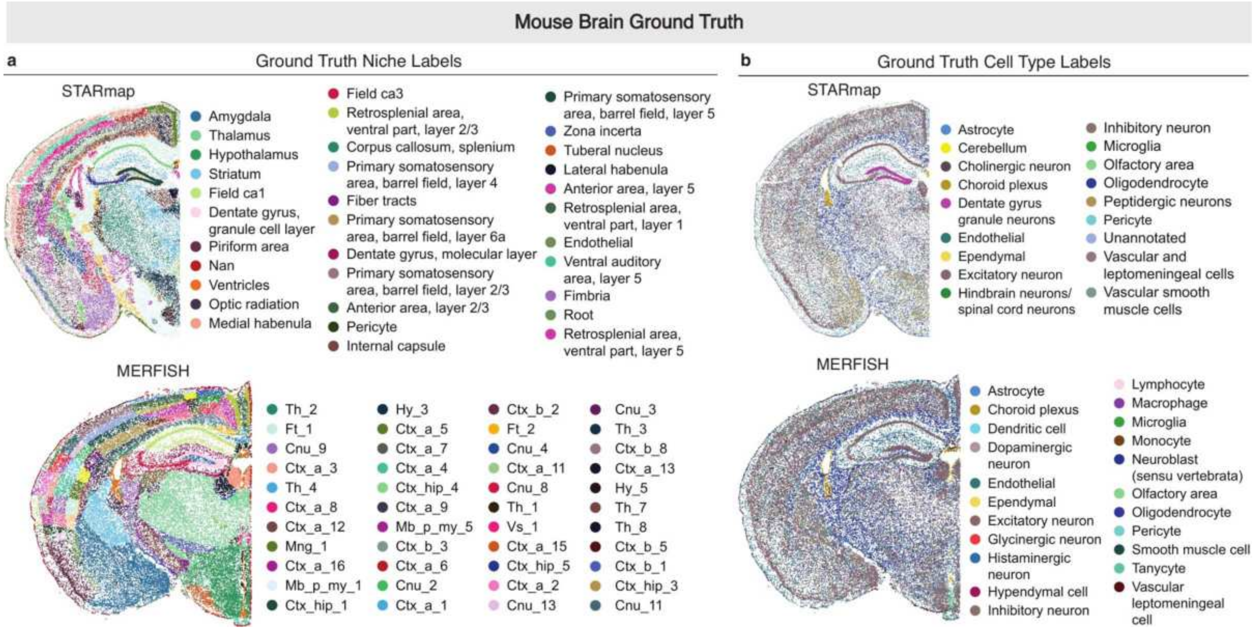


Fig. S2: Mouse-brain (STARmap + MERFISH) expert ground-truth annotations. **(a)** Ground-truth niche labels and **(b)** ground-truth cell-type labels on the STARmap (top) and MERFISH (bottom) sections, against which the Mouse-Brain NMI / ARI in Table 1 are computed.

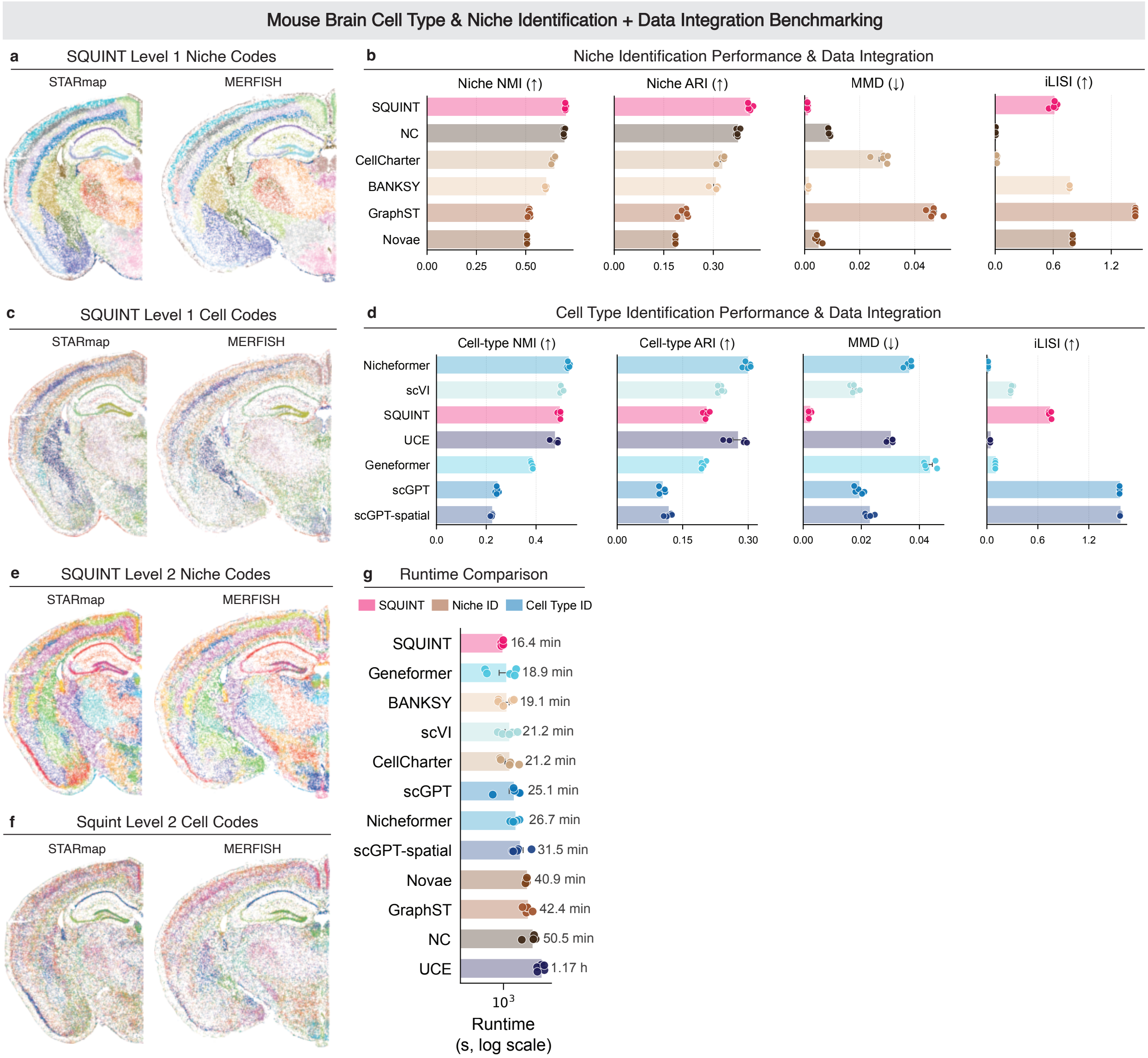


Fig. S3: Extended niche- and cell-type-identification and integration benchmark on the mouse brain (STARmap + MERFISH). **(a, c)** SQUINT Level-1 niche (a) and cell (c) codes, and **(e, f)** the finer Level-2 niche (e) and cell (f) codes, on both sections. **(b, d)** Niche-branch (b) and cell-branch (d) identification (NMI, ARI) and integration (MMD, iLISI): GraphST (b) and scGPT / scGPT-spatial (d) reach high iLISI but weak NMI / ARI (over-integration), whereas SQUINT pairs a moderate iLISI with the lowest MMD (Sections 4.2 and 4.3). **(g)** Wall-clock runtime (log scale). Bars: mean over five seeds; dots: seeds.

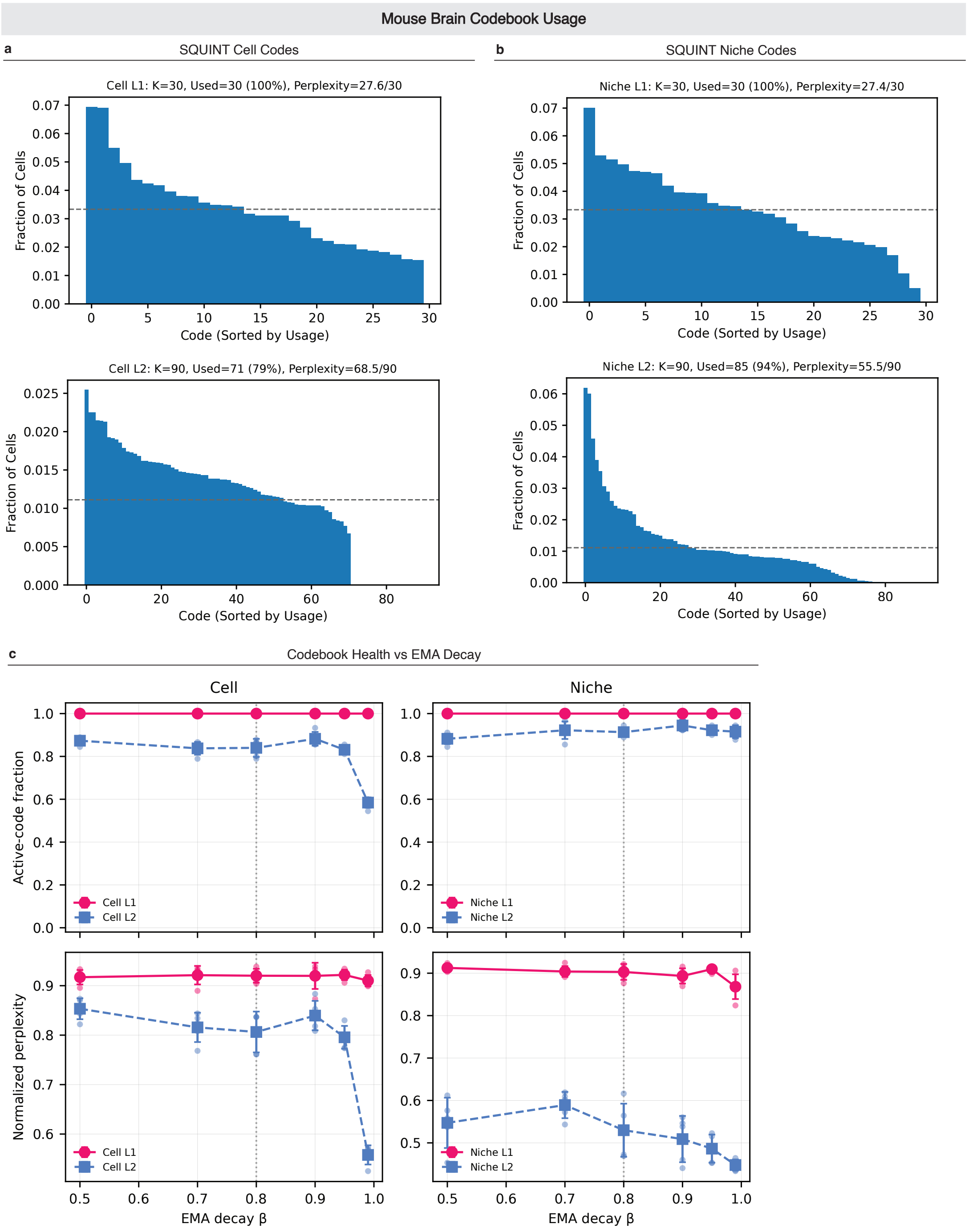


Fig. S4: Codebook usage and EMA-decay sensitivity on the paired mouse-brain STARmap + MERFISH benchmark. **(a, b)** Fraction of cells assigned to each code, sorted by usage, for the **(a)** cell and **(b)** niche codebooks at the coarse (Level-1, $K^{(1)}$=30) and fine (Level-2, $K^{(2)}$=90) residual levels (dashed line: uniform usage; per-panel active fraction and perplexity in the titles) — codes are used densely rather than collapsing onto a few entries. **(c)** Codebook health — active-code fraction and normalized perplexity (mean over five seeds; per-seed dots) — versus the EMA decay $\beta$ for the cell and niche codebooks; the dotted line marks our default $\beta$=0.8. Health is stable across $\beta \in [0.5, 0.95]$, with Level-1 fully used throughout; only the extreme $\beta$=0.99 starves the fine cell codebook, so $\beta$=0.8 sits in a healthy plateau. Cosine-similarity VQ with EMA updates and dead-code resampling (Appendix A) avoids codebook collapse.

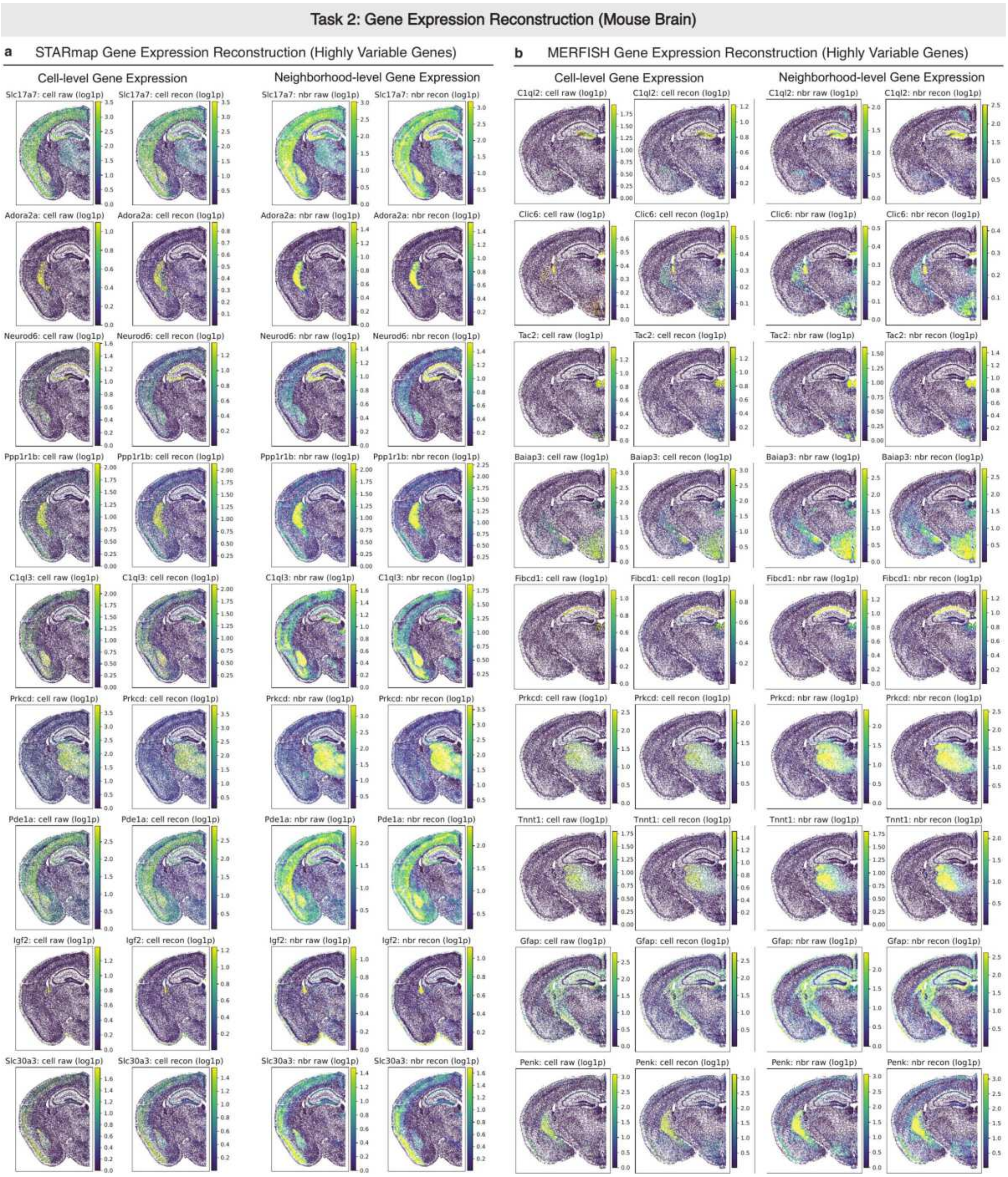


Fig. S5: Extended Task 2 (gene-expression reconstruction; mouse-brain STARmap + MERFISH). Reconstructions of highly-variable genes on **(a)** STARmap and **(b)** MERFISH, raw (left) versus SQUINT (right), at the cell and 1-hop neighbourhood level.

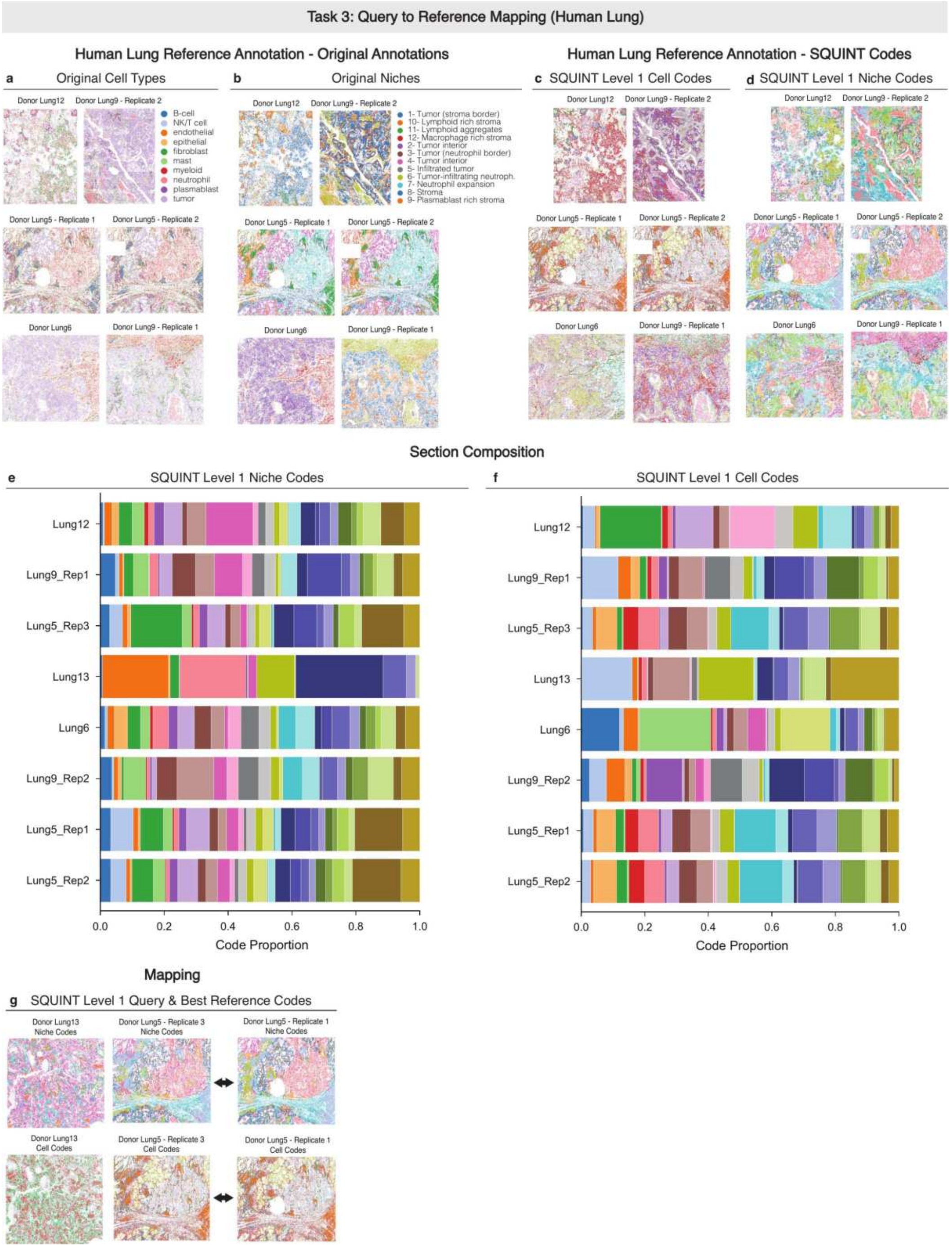


Fig. S6: Extended Task 3 (query-to-reference mapping; CosMx human NSCLC). **(a, b)** Expert cell-type (a) and niche (b) annotations on the reference and query sections. **(c, d)** SQUINT Level-1 niche (c) and cell (d) codes on the same sections, recapitulating the expert organisation across donors and replicates. **(e, f)** Per-section composition of SQUINT Level-1 niche (e) and cell (f) codes, the code distributions underlying the similarity in Fig. 3a; sections sharing a donor have visibly similar compositions. **(g)** Each query's Level-1 niche and cell codes beside its best-matching reference section.

Table S1: **Capacity ablations.** Encoder and codebook-size axes (mouse-brain STARmap + MERFISH); conventions as in Table 3. Codebook blocks sweep one RVQ level ($K^{(1)}$, $K^{(2)}$) with the other fixed (shaded = sweep anchor; global default $(30, 90)$). Neighbours: 16/8 = 16-graph with 8 sampled. Defaults are chosen for balance across metrics and cost, so a sweep setting can marginally edge one metric (e.g. $k$=24 neighbours, niche NMI 0.710 vs. 0.702) without warranting the change.

| | **GNN depth** | | **Neighbours** | | | | **Cell cb.** $K^{(1)}$ | | | | **Cell cb.** $K^{(2)}$ | | | | **Niche cb.** $K^{(1)}$ | | | | **Niche cb.** $K^{(2)}$ | | | |
|---|---|---|---|---|---|---|---|---|---|---|---|---|---|---|---|---|---|---|---|---|---|---|
| *Variant* | 1L | 2L | 16 | 8 | 16/8 | 24 | (30,30) | (10,30) | (90,30) | (300,30) | (30, 90) | (30,10) | (30,30) | (30,300) | (30,30) | (10,30) | (90,30) | (300,30) | (30, 90) | (30,10) | (30,30) | (30,300) |
| Niche NMI↑ | 0.702 | $0.675^{**}$ | 0.702 | $0.661^{***}$ | $0.685^{**}$ | $0.710^{**}$ | 0.702 | 0.703 | 0.702 | 0.701 | 0.702 | 0.704 | 0.702 | 0.702 | 0.699 | $0.600^{***}$ | $0.689^{**}$ | 0.701 | 0.702 | 0.701 | 0.699 | 0.703 |
| ARI↑ | 0.410 | $0.379^{**}$ | 0.410 | $0.383^{***}$ | 0.404 | 0.412 | 0.410 | 0.414 | 0.414 | 0.413 | 0.410 | 0.412 | 0.410 | 0.410 | 0.404 | $0.246^{***}$ | $0.360^{***}$ | 0.410 | 0.410 | 0.411 | 0.404 | 0.412 |
| MMD↓ | 1.05 | 1.52 | 1.05 | 0.97 | 1.09 | 1.11 | 1.01 | 1.13 | $1.46^{**}$ | $2.53^{***}$ | 1.05 | 1.02 | 1.01 | 0.99 | 0.98 | 0.96 | 1.10 | 1.02 | 1.05 | 0.96 | 0.98 | 1.03 |
| iLISI↑ | 0.609 | 0.551 | 0.609 | 0.651 | 0.598 | 0.589 | 0.634 | $0.574^{*}$ | 0.605 | $0.558^{**}$ | 0.609 | 0.627 | 0.634 | 0.620 | 0.623 | 0.572 | 0.636 | 0.617 | 0.609 | 0.584 | 0.623 | 0.572 |
| Cell NMI↑ | 0.494 | 0.491 | 0.494 | 0.490 | 0.494 | 0.493 | 0.490 | $0.446^{***}$ | $0.458^{***}$ | $0.434^{***}$ | 0.494 | 0.490 | 0.490 | 0.493 | 0.490 | 0.492 | 0.493 | $0.495^{*}$ | 0.494 | 0.491 | 0.490 | 0.490 |
| ARI↑ | 0.204 | 0.201 | 0.204 | 0.198 | 0.205 | 0.201 | 0.200 | $0.268^{***}$ | $0.097^{***}$ | $0.040^{***}$ | 0.204 | 0.201 | 0.200 | 0.201 | 0.200 | 0.203 | 0.203 | $0.209^{*}$ | 0.204 | 0.200 | 0.200 | 0.203 |
| MMD↓ | 2.27 | 2.51 | 2.27 | 2.44 | 2.20 | 2.34 | 2.12 | $1.73^{*}$ | 2.52 | $3.04^{**}$ | 2.27 | $1.64^{*}$ | 2.12 | 2.43 | 2.43 | 2.26 | $2.02^{*}$ | 2.53 | 2.27 | 2.19 | 2.43 | 2.49 |
| iLISI↑ | 0.739 | 0.737 | 0.739 | 0.734 | 0.745 | 0.751 | 0.709 | $0.569^{**}$ | 0.734 | $0.640^{**}$ | 0.739 | $0.614^{***}$ | 0.709 | 0.737 | 0.751 | 0.747 | 0.739 | 0.741 | 0.739 | 0.755 | 0.751 | 0.737 |

#### Appendix E. Discussion

**Towards self-supervised transformer pretraining on tissues.** A central advantage of a *discrete* representation is that it turns each tissue into a sequence of tokens that can be consumed directly by transformer-style self-supervised objectives (masked- or next-token prediction, contrastive sequence modelling) without the bespoke discretisation step that current spatial foundation models rely on (gene-rank tokenisation in Nicheformer [16], $k$-means in GeST [7]). Because SQUINT's tokens already encode both cell-intrinsic and microenvironment identity, the resulting tissue sequences carry the structure a foundation model would otherwise have to recover from raw counts, a concrete path to tissue-scale pretraining and zero-shot transfer to new microenvironments.

**Limitations and future work.** SQUINT's cell codebook is already competitive with foundation-model baselines on cell-type identification and cell-level imputation (Sections 4.3 and 4.4); it could be further improved by deeper RVQ stacks, by *joint cell–niche decoding* that conditions the cell decoder on both $\mathbf{z}^{q,\text{cell}}$ and $\mathbf{z}^{q,\text{niche}}$, or by initialising the cell branch from a single-cell foundation model and fine-tuning end-to-end. Our cross-assay integration result rests on a single STARmap + MERFISH pair – the other cohorts are single-assay – so establishing cross-assay alignment as a general property will require further assay pairings. Expanding the benchmark to more datasets (additional SRT assays, tissues, donors and disease contexts) is thus the natural next step and a prerequisite for using SQUINT as a tokeniser for atlas-scale foundation models. Multi-modal extensions (histology, protein measurements, cross-organ transfer) build on the same architecture.

[3] Huiwen Chang, Han Zhang, Lu Jiang, Ce Liu, and William T. Freeman. MaskGIT: Masked generative image transformer. In *Proceedings of the IEEE/CVF Conference on Computer Vision and Pattern Recognition (CVPR)*, 2022.
